# Mapping fetal brain development based on automated segmentation and 4D brain atlasing

**DOI:** 10.1101/2020.05.10.085381

**Authors:** Haotian Li, Guohui Yan, Wanrong Luo, Tintin Liu, Yan Wang, Ruibin Liu, Weihao Zheng, Yi Zhang, Kui Li, Li Zhao, Catherine Limperopoulos, Yu Zou, Dan Wu

## Abstract

Fetal brain MRI has become an important tool for in utero assessment of brain development and disorders. However, quantitative analysis of fetal brain MRI remains difficult, partially due to the limited tools for automated preprocessing and the lack of normative brain templates. In this paper, we proposed an automated pipeline for fetal brain extraction, super-resolution reconstruction, and fetal brain atlasing to quantitatively map in utero fetal brain development during mid-to-late gestation in a Chinese population. First, we designed a U-net convolutional neural network for automated fetal brain extraction, which achieved an average accuracy of 97%. We then generated a developing fetal brain atlas, using an iterative linear and nonlinear registration approach. Based on the 4D spatiotemporal atlas, we quantified the morphological development of the fetal brain between 23-36 weeks of gestation. The proposed pipeline enabled the fully-automated volumetric reconstruction for clinically available fetal brain MRI data, and the 4D fetal brain atlas provided normative templates for quantitative analysis of potential fetal brain abnormalities, especially in the Chinese population.

## Introduction

Magnetic resonance imaging (MRI) provides superior and versatile contrasts of the gray and white matter structures in the fetal brains without known safety concerns, and its role in fetal brain examination has been recognized over the years (Griffiths et al., 2017; Jarvis & Griffiths, 2019; Weisstanner, Kasprian, Gruber, Brugger, & Prayer, 2015). In utero MRI offers exquisite anatomical details of the fetal brain within millimeter resolution and has become an important tool for prenatal diagnosis, complementary to ultrasound examination (Nielsen & Scott, 2017). For instance, gyral and sulcal abnormalities (Rolo et al., 2011), corpus callosum dysgenesis (Glenn, 2006), and abnormal cortical maturation (Fogliarini et al., 2005) in the fetal brain could be revealed by in utero MRI. Moreover, quantitative analysis of in utero images further improved our understanding of fetal brain development (Makropoulos, Counsell, & Rueckert, 2018; Scott et al., 2011).

Three-dimensional (3D) high-resolution images are often required for quantitative analysis of the brain, which, however, remains challenging for in utero MRI, due to the excessive and unpredictable fetal and maternal abdominal motion. Slice-to-volume registration (SVR) (Jiang et al., 2007; Kainz et al., 2015; Francois Rousseau et al., 2006) of 2D multi-slice images with the super-resolution (SR) technique (Gholipour, Estroff, & Warfield, 2010; Kuklisova-Murgasova, Quaghebeur, Rutherford, Hajnal, & Schnabel, 2012; François Rousseau, Kim, Studholme, Koob, & Dietemann, 2010) is commonly used to obtain volumetric reconstruction of the fetal brains. To achieve accurate SR reconstruction, extraction of the fetal brain from in utero images is required, which relies on the manual delineation of the brain contours on 2D slices in all three orientations. This labor-intensive process inhibits large data analysis.

Several brain extraction methods have been proposed and are widely available in published toolboxes, including the Brain Extraction Tool (BET) in FSL (Jenkinson, Pechaud, & Smith, 2005; Smith, 2002), 3dSkullStrip in the AFNI toolkit (Cox, 1996), the Hybrid Watershed Algorithm (HWA) in FreeSurfer (Lin et al., 2003), and Robust Learning-Based Brain Extraction (ROBEX) (Iglesias, Liu, Thompson, & Tu, 2011). However, these brain extraction methods developed for the adult brain usually fail for the fetal brain, due to the complex in utero and abdominal tissues surrounding the fetal brain. Deep learning based segmentation methods have been proposed in recent years. Kleesiek et al. used a convolutional neural network (CNN) for adult brain extraction, with used cubic windows of a fixed size around each voxel (Kleesiek et al., 2016). Salehi et al. proposed an auto-context CNN (auto-net) with two 2.5D network architectures and developed a geometry-independent and registration-free adult brain extraction tool (Salehi, Erdogmus, & Gholipour, 2017). These methods have also been extended to segment the fetal brain, which achieved higher segmentation accuracy compared to the traditional approaches. Therefore, we proposed a model based U-net (Ronneberger, Fischer, & Brox, 2015) to directly segment the fetal brain from routine clinical in utero MRI data acquired in three orientations, as a preprocessing step before SR reconstruction.

Another challenge for in utero fetal brain MRI is that the rapid developmental changes of the fetal brain impose difficulties for radiological examinations that mainly rely on visual inspection and empirical assessment. It is essential to have normative fetal brain templates at matching gestational ages (GA) to compare with. Brain templates or atlases play an important role in quantitative image analysis. Currently, the development of fetal brain atlases is limited compared with that of neonatal, pediatric and adult brain atlases, due to the difficulties in the acquisition and preprocessing. Initial attempts have been made. Habas et al. developed a probabilistic fetal brain MRI atlas using clinical MR scans of 20 young fetuses with GA ranging from 20.6 to 24.7 weeks (Habas et al., 2010). Serag et al. constructed a 4D atlas of the developing fetal brains between 23 and 37 weeks of gestational, using T2 weighted MR images from 80 fetuses (Ahmed Serag et al., 2012; A Serag et al., 2012) (https://brain-development.org/brain-atlases/fetal-brain-atlases/fetal-brain-atlas-serag/). Gholipour et al. established an unbiased four-dimensional atlas of the fetal brain using high-resolution MRI of 81 normal fetuses between 19 and 39 weeks of gestation (Gholipour et al., 2017) and made it a public resource (http://crl.med.harvard.edu/research/fetal_brain_atlas/). However, all the aforementioned fetal brain atlases were established in Caucasian populations. It is known that there are considerable anatomical and functional differences between Caucasian and Asian cohorts in the pediatric, adolescent, and adult brains (Lee et al., 2005; Tang et al., 2010; Uchiyama, Seki, Tanaka, & Koeda, 2013). It is likely that these differences start in the fetal period due to genetic factors (Rao et al., 2017). Therefore, the existing atlases may not be ideal for the analysis of fetal brains in a non-Caucasian population. Given the rapid development of the fetal brain, a small difference between the subject and atlas may have a noticeable impact. Here, we generated the first version of Chinese fetal brain atlas between 23-36 weeks of gestation, which allowed us to quantitatively characterize the three-dimensional morphological evolution of the fetal brain.

## Methods

### 1. Dataset

In our study, all the data were collected retrospectively from routine clinical scans at Women’s Hospital of Zhejiang University School of Medicine between the years of 2013-2019. The research protocols were approved by the local Institutional Review Board with a waiver of consent. In utero MRI images from pregnant women between 21 to 40 weeks of pregnancy were included in this study. Exclusion criteria for the normal pregnancy included suspected fetal growth restriction based on ultrasound screening, fetal intracranial abnormalities such as ventriculomegaly and cerebral hemorrhage, chromosome abnormalities, gestational diabetes mellitus, and maternal intrauterine infections including cytomegalovirus and toxoplasmosis.

The scans were performed at a 1.5 T GE scanner (Signa Hdxt) with an 8-channel cardiac coil. No sedation or contrast agents were administered in this study. Images were acquired using the single shot Fast Spin Echo (ssFSE) or the T2-prepared balanced Steady State Free Precession (SSFP) sequence. The ssFSE data was acquired with repetition time (TR) = 2400 ms, echo time (TE) = 130 ms, field-of-view (FOV) = 360×360 mm, imaging matrix = 512×512 (in-place resolution = 0.7×0.7 mm), and approximately 20 slices with slice thickness of 4±0.1 mm and no slice gap. The bSSFP data was acquired at TR = 4.7 ms, TE = 2.1 ms, flip angle = 55°, FOV = 380×380 mm, imaging matrix = 512×512 (in-place resolution = 0.74×0.74 mm), and approximately 16 slices with slice thickness of 5±0.1 mm and no slice gap.

In total, we obtained 636 scans from 212 fetal brains in axial, coronal, and sagittal orientations after visual inspection for image quality. The distribution of data at each GA is shown in Figure 1, separately for the two types of sequences in three orientations. Note that the three orientations of the same fetus may be scanned with different sequences. The SSFSE and bSSFP images in this study had comparable contrasts, and therefore, they were jointly used for the U-net based brain segmentation.

**Figure 1:**
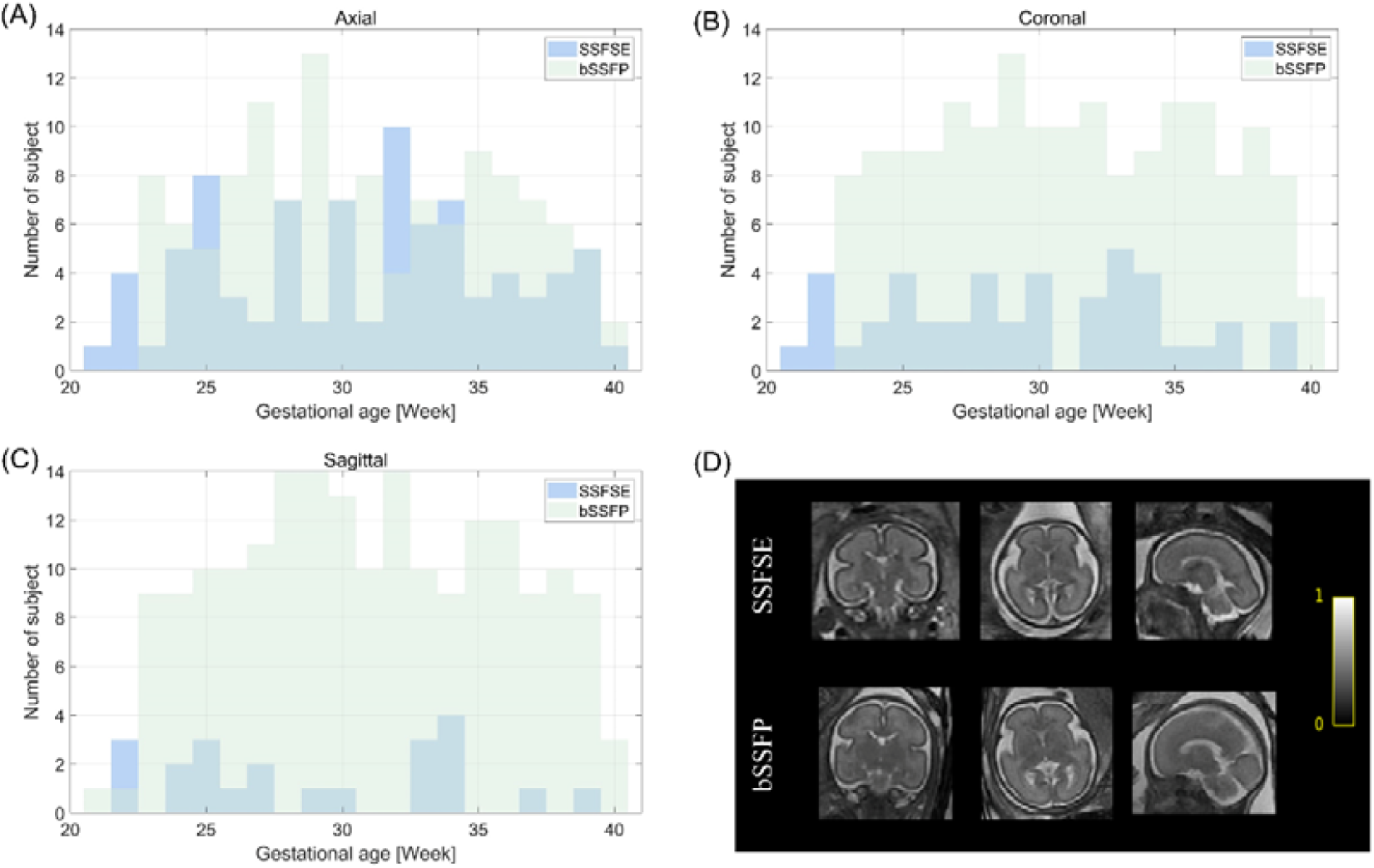
Distribution of the fetal brain MRI data used for the U-net based fetal brain extraction. The images were acquired in the axial (A), coronal (B), and sagittal (C) orientations between 21-40 gestational weeks, by SSFSE and bSSFP sequences. Two fetal brain images at 26 weeks of gestation in three orientations using the SSFSE and the bSSFP sequences respectively are indicated in the lower right corner. (D) Fetal brain images acquired using the SSFSE sequence and bSSFP sequence.

47 fetal brains were diagnosed as radiologically and clinically normal by experienced radiologists and clinicians (ZY, YG, and LK). After removing 12 normal fetal brains that had noticeable motion artifacts or low SNR, 35 normally developing fetal brains between 23-36 weeks of gestation were selected (five brains every two gestational weeks) for atlas generation. The 35 brains were acquired with the bSSFP sequence in three orientations.

### 2. Fetal brain extraction

#### 2.1 2D U-Net architecture

The fetal brain masks were manually delineated by a trained research assistant on all 636 scans and used as the ground truth for training in the following network. Manual brain extraction took about thirty to forty minutes per fetal brain (including three orientations), depending on the GA of the fetal brain and the quality of the images.

Figure 2 shows the proposed U-Net CNN structure for fetal brain segmentation. The U-net has an approximately symmetric structure, which consists of a contracting path and an expanding path. Each convolutional layer is followed by a ReLU non-linear layer. In the contracting path, a 2×2 max-pooling layer is applied after two 3×3 convolutional layers and the number of feature channels gets doubled. Correspondingly, the expanding path utilizes a 2×2 up-sampling layer after a convolutional operation to halve the feature channels. A dropout rate of 0.5 is used before the last pooling layer and the first up-sampling layer. The sizes of the symmetric structures between the contracting and the expanding paths are kept the same. Therefore, the outputs of the contracting path are directly concatenated with the corresponding layers of the expanding path. In the final layer, a 3×3 convolutional layer converts the feature maps to label probability, and a 1×1 convolution layer with linear output predicts the fetal brain contour.

**Figure 2:**
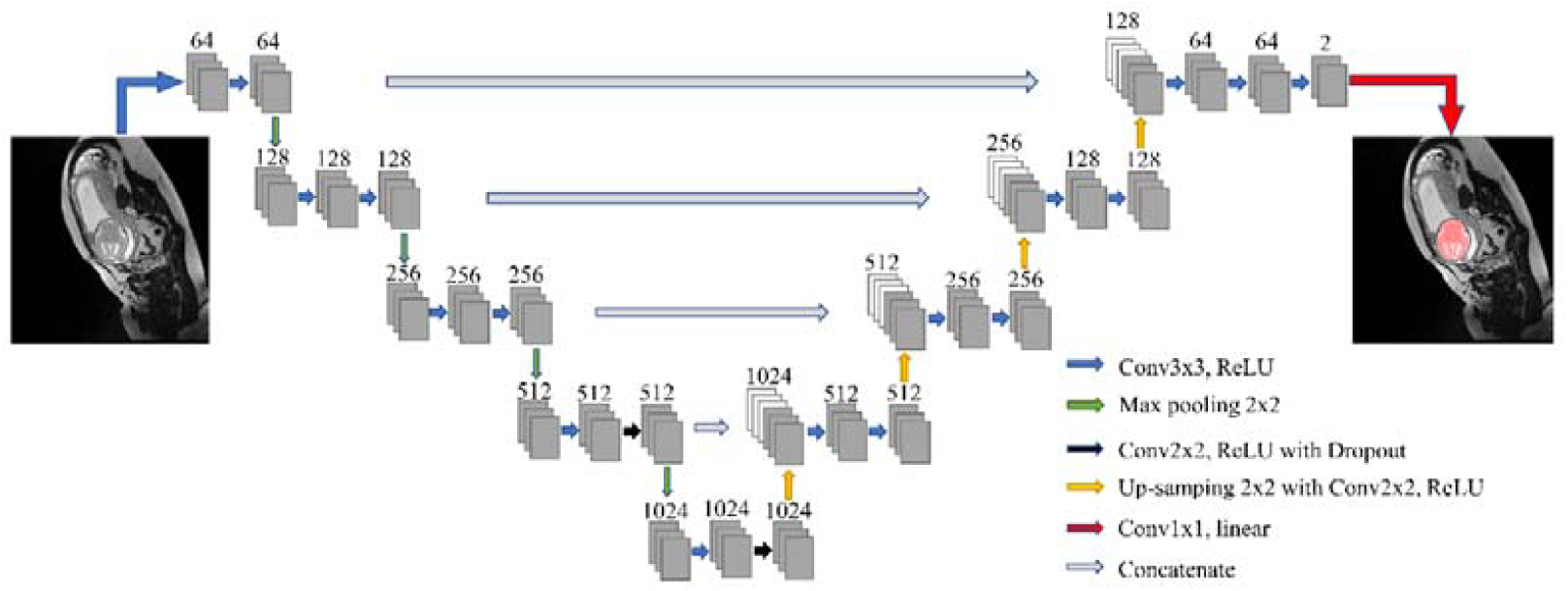
The U-net convolutional network for fetal brain segmentation. The network consisted of a contraction path and an expansion path. The convolution layer was set to have a kernel size of 3×3 and stride of 2 with zero-padding. The number of features is labeled above the network layers, and the types of connection between layers are indicated in the lower right corner.

Images acquired in three orientations were separately trained using independent U-net, which shared the same network structure. There were 212 scans from 212 fetal brains for each network, and we used individual slices as input data to the U-nets. The data were randomly divided into 6:2:2 for training, validation, and testing within each gestational week. Cross-validation was used to make the best use of the data and check potential overfitting. Image data in the training and validation sets were enhanced by ten times with image translation ranged 0-20 pixels, rotation ranged 0-20 degrees, random cropping, and vertical mirror symmetry (Perez & Wang, 2017).

We applied the Softmax method to measure the loss function for every pixel, and cross-entropy loss function between predicted results and ground truth was minimized on 30 epochs. The U-net network was trained using an ADAM optimizer with an initial learning rate of 1×10^−6^ that was multiplied by 0.1 for every 10 epochs. The training time was approximately six hours with four-parallel Nvidia Geforce GTX2080Ti GPUs. For testing, we obtained a probability image as the output of the network, and the final mask was determined using a threshold of 0.9. The choice of threshold (between 0.1-0.9) did not affect the segmentation accuracy.

### 2.2 Evaluation of the segmentation accuracy

The test set of fetal brain images was also segmented using the brain extraction tool (BET) (Jenkinson et al., 2005), which uses a deformable spherical surface mesh model initialized at the center-of-gravity of the image. The input images for BET were cropped from 512×512 to 256×256 to reduce the influence of surrounding tissues as the fetal brains laid in the center of the images. The neighborhood filling was performed for both the U-net and the BET outputs to remove the holes and islands. The performance of the U-Net and BET methods was assessed by comparison with the manual brain segmentation based on the Dice score, intersection over union (IOU), sensitivity, and specificity in three orientations. Based on the predicted brain mask A and the ground truth mask B, the true positive (TP), false positive (FP), true negative (TN) and false negative (FN) rates were calculated, and Dice is defined as 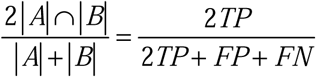, IOU as 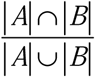, specificity as 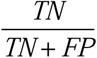, and sensitivity as 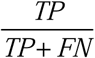.

### 3. Super-resolution (SR) reconstruction

3D fetal brain images were reconstructed using a SR pipeline (Francois Rousseau et al., 2006) based on the 2D fetal brain images extracted in three orientations. We used an open-source toolkit “BTK” (François Rousseau et al., 2013) (https://github.com/rousseau/fbrain) to perform global histogram matching among the axial, sagittal, and coronal images, non-local denoising, SVR registration, and SR reconstruction. We used the default parameters except for the regularization factor of the SR algorithm which was increased for higher contrast.

### 4. Generation of the fetal brain atlas

We designed an iterative linear and nonlinear registration framework to construct the fetal brain atlas based on SR reconstructed 3D fetal brains. We selected five normal brains from every two gestational week bins to generate population templates. The atlas generation pipeline is shown in Figure 3. Among the five brains in each bin, we carried out pairwise registrations (Ahmed Serag et al., 2012) by selecting one of the brains as a target image and the rest of the brains were registered to the target brain, by affine and Symmetric image Normalization (SyN) registration (Avants, Epstein, Grossman, & Gee, 2008), using the ANTs toolbox (https://github.com/ANTsX/ANTs). This procedure was applied to each of the five brains and produced a group average for every brain. Averaging the five groups averaged images generated the initial template (*I*_*A*_^1^). The use of pairwise registrations eliminated bias in the atlas toward any of the original images. In the first iteration, the five brains were registered to the *I*_*A*_^1^ from their native space using affine and deformable SyN registration, and averaged to obtain an averaged template (*I*_*A*_^2^). In the second iteration, all brains were transformed to *I*_*A*_^2^ from their native space with affine and deformable SyN registration, and averaged to get an averaged template (*I*_*A*_^3^). The procedure was repeated 15 times until the template became stable (Supplementary Figure 1).

**Figure 3:**
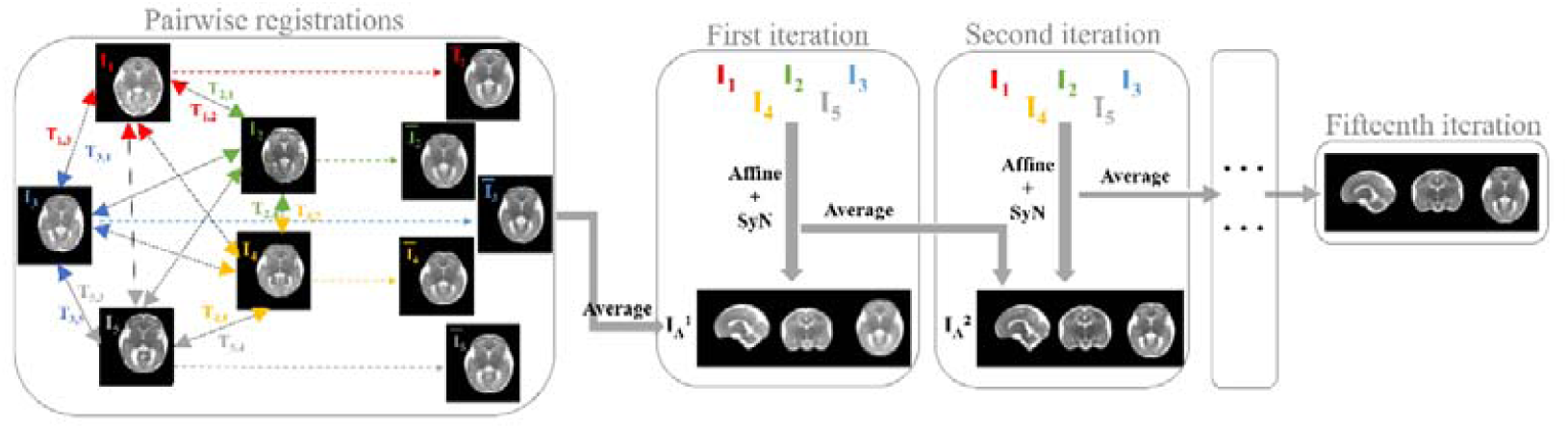
Pipeline for fetal brain atlas generation. Five normal developing fetal brain images were chosen at a given gestational stage (every two weeks). Pairwise registrations using affine and SyN registration were performed to generate the initial template I_A_^1^. In the first iteration, the five fetal brains were registered to the I_A_^1^ from their native space by affine and SyN transformation to generate I_A_^2^. The procedure was iterated 15 times to produce the final template.

### 5. Mapping the morphological fetal brain development

To quantify the morphological changes of the fetal brain parenchyma, we removed the cerebrospinal fluid (CSF) on the fetal brain atlas. The atlas images were first segmented by the Developing Brain Region Annotation with Expectation-Maximization (Draw-EM) tool (Makropoulos et al., 2014). The segmentation results were then manually corrected by a trained research assistant in ROIEditor (https://www.mristudio.org/).

Based on the CSF-free fetal brain atlas, morphological changes between gestational stages were quantified by the transformation between adjacent gestational stages. For instance, transforming the template at 23-24 weeks to the template at 25-26 weeks to obtain the morphological change from 23.5 weeks to 25.5 weeks. The transformation was achieved by rigid registration followed by SyN registration. For each pair of transformation, the deformation field was computed using the log-Jacobian matrix of the SyN transformation. The determinant of the log-Jacobian matrix was used to quantify the amount of morphological differences between adjacent gestational stages. In addition, the dynamic changes from 23 to 36 weeks of gestation can be captured continuously by interpolating the log-Jacobian matrices, and predicted brain atlas can be obtained at any given GA.

## Results

Figure 4 shows representative segmentation results of fetal brains at different GAs in the sagittal, coronal, and axial orientations. The automated segmentation by the U-net (magenta contours), mostly overlapped with the manually delineated ground truth (green contours). Table 1 demonstrates the segmentation performance of the U-net method compared with the BET method in the test set, in three orientations. The U-net method yielded an average Dice score of 0.97 across the three brain orientations (0.9774, 0.9759, and 0.9564 in the coronal, sagittal, and axial orientations, respectively). In comparison, the BET method resulted in an average Dice score of 0.74 and significantly lower IOU, sensitivity, and specificity in all three orientations.

**Table 1:** The performance of the U-net model and conventional BET method. The Dice score, IOU, specificity, and sensitivity of the segmentation results in three orientations were compared between the two methods.

|  | *Dice* |  |  | *IOU* |  |  |
| --- | --- | --- | --- | --- | --- | --- |
| Methods | Coronal | Sagittal | Axial | Coronal | Sagittal | Axial |
| U-net | 0.9774 ± | 0.9759 ± | 0.9564 ± | 0.9558 ± | 0.9531% ± | 0.9170 ± |
|  | 0.0074 | 0.0121 | 0.0176 | 0.0139 | 0.0226 | 0.0320 |
| BET | 0.7406 ± | 0.7284 ± | 0.7458 ± | 0.6668 ± | 0.6731 ± | 0.6550 ± |
|  | 0.3226 | 0.3686 | 0.2737 | 0.3192 | 0.3565 | 0.2875 |
|  | *Specificity* |  |  | *Sensitivity* |  |  |
| Methods | Coronal | Sagittal | Axial | Coronal | Sagittal | Axial |
| U-net | 0.9994 ± | 0.9992 ± | 0.9982 ± | 0.9762 ± | 0.9744 ± | 0.9723 ± |
|  | 0.0003 | 0.0004 | 0.0009 | 0.0109 | 0.0216 | 0.0128 |
| BET | 0.9848 ± | 0.9892 ± | 0.9828 ± | 0.8444 ± | 0.7404 ± | 0.8823 ± |
|  | 0.0204 | 0.0164 | 0.0191 | 0.3380 | 0.3723 | 0.2659 |

**Figure 4:**
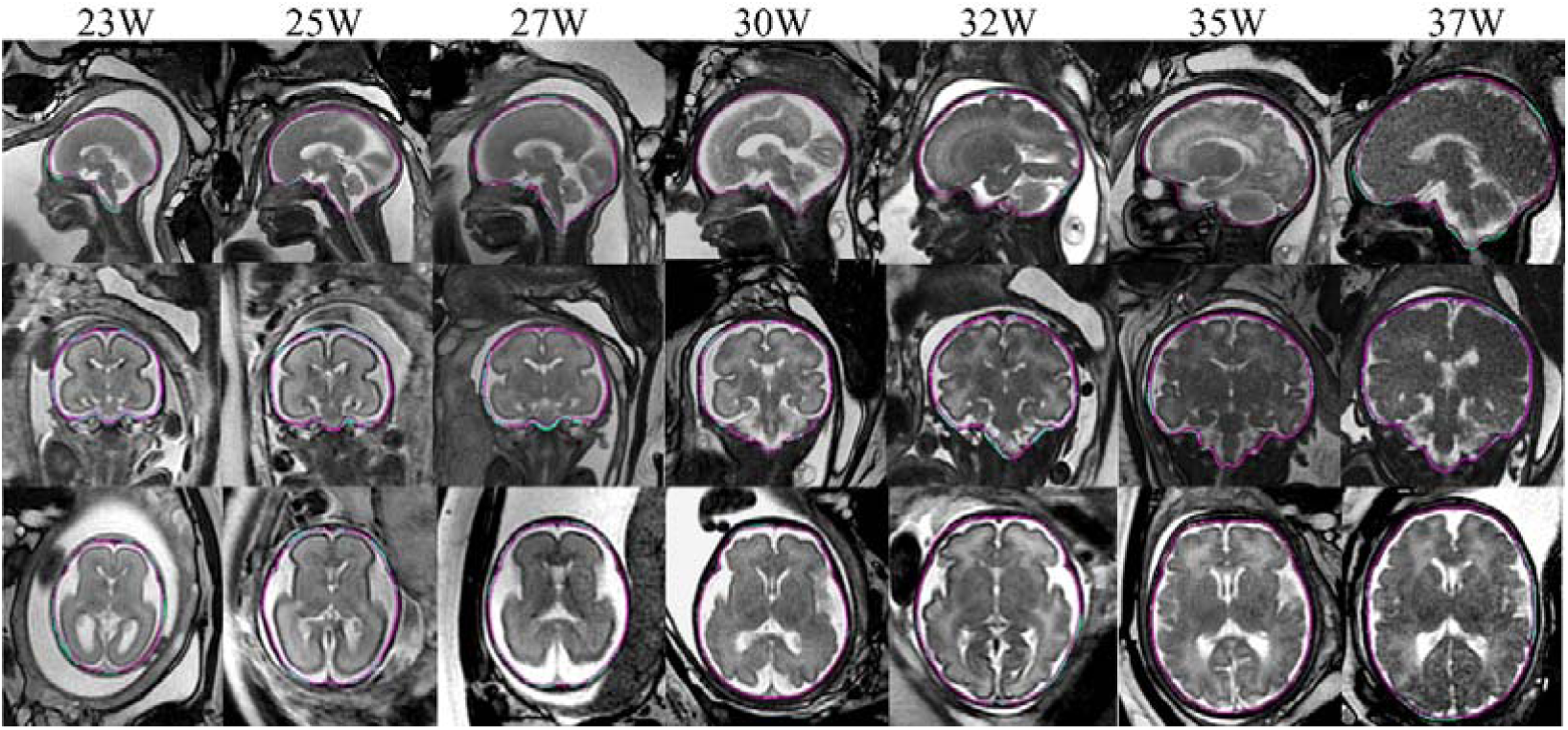
The contours of the U-net predicted brain mask (in magenta) and the ground truth (in green) are shown for fetal brains at different gestational weeks, in sagittal, coronal, and axial orientations.

We observed a GA dependent variation of the segmentation accuracy in our results (Figure 5), especially in the axial images. The relatively low accuracy at early GA was likely to be related to the relatively small number of training data and relatively small brain size compared with the majority of the data. Besides, Dice scores in the coronal and sagittal orientations were higher than those in the axial orientation, which was possibly due to the difficulty in segmenting bottom part of the fetal brain in the axial orientation, e.g., the medulla oblongata and the cerebellum. Nevertheless, the overall high segmentation accuracy was sufficient for SR reconstruction.

**Figure 5:**
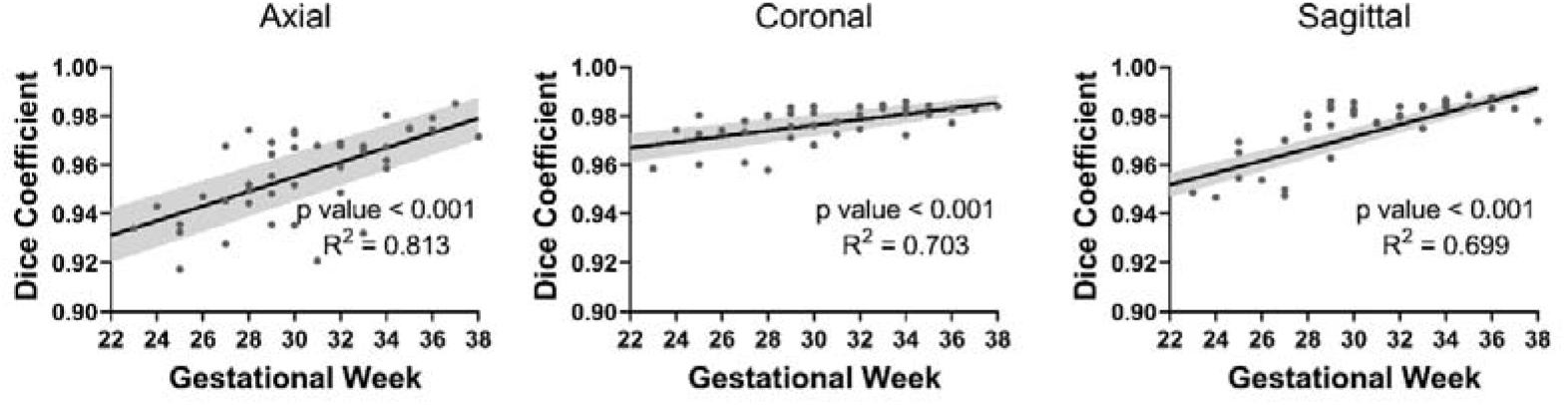
Relation between the Dice scores from the U-net segmentation and GA in three orientations. The solid line indicates the mean and the shaded region indicates the standard deviation of the Dice scores at varying gestational weeks.

Based on the U-net masked images, we constructed the 3D fetal brain images using SR reconstruction (Supplementary Figure 2). We then constructed the fetal brain atlas by computing the average brain templates every two weeks from 23 to 36 gestational weeks. The 4D spatiotemporal fetal brain atlas is shown in Figure 6, in sagittal, coronal, and axial views, which characterized the drastic changes in the shape and size of the fetal brains. It was observed that between 23 to 26 weeks of gestation, the development of calcarine fissure (Bendersky, Musolino, Rugilo, Schuster, & Sica, 2006) and cingulate gyrus (Monteagudo & Timor-Tritsch, 1997) were prominent; part of the primary sulci, precentral gyrus, and postcentral gyrus were formed between 27 to 30 weeks of gestation; the rest of the primary sulci and part of the secondary sulci appeared between 31 to 34 weeks of gestation; and between 34 to 36 weeks of gestation, the ventricles gradually shrink due to the expansion of brain parenchyma (Huisman, Martin, Kubik-Huch, & Marincek, 2002). In addition, to fill the gestational gap (2 weeks) in the current atlas, we interpolated the transformation matrices between adjacent GA to generate pseudo-templates at a 0.5-week interval (Supplementary Figure 3), or even finer intervals (Supplementary video) for better visualization of the dynamic process.

**Figure 6:**
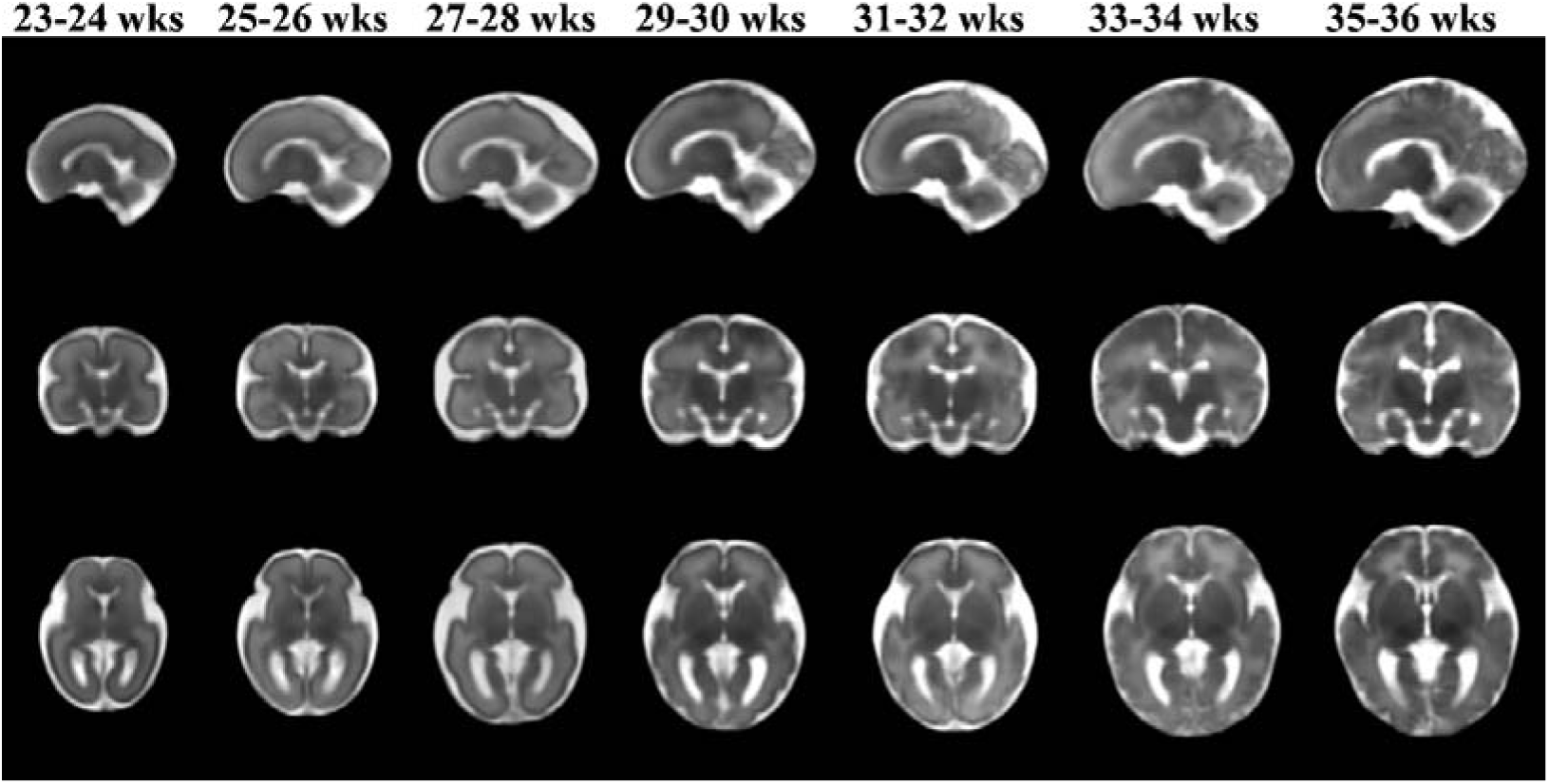
Fetal brain atlas was generated for GAs of 23-24, 25-26, 27-28, 29-30, 31-32, 33-34, and 35-36 weeks, in sagittal, coronal, and axial views.

The morphological changes between adjacent gestational stages are illustrated in Figure 7 in 2D and 3D views. The color bar represents the amount of morphological deformation when transforming one template to the next, indicating the rate of brain growth from one gestational stage to the next. Dramatic fetal brain growth was observed during early gestation, e.g., from 23.5 to 25.5 weeks, and the growth rate slowed down towards late gestation. Moreover, a posterior-to-anterior developing pattern was observed. The log-Jacobian map of brain transformation from 27.5 to 29.5 weeks indicated the fastest changes in the central and posterior brain including the central sulcus, pre- and post-central gyri, and occipital lobe regions (white arrows), while the transformation from 31.5 to 33.5 weeks suggested prominent changes in the frontal-orbital regions (blue arrows).

**Figure 7:**
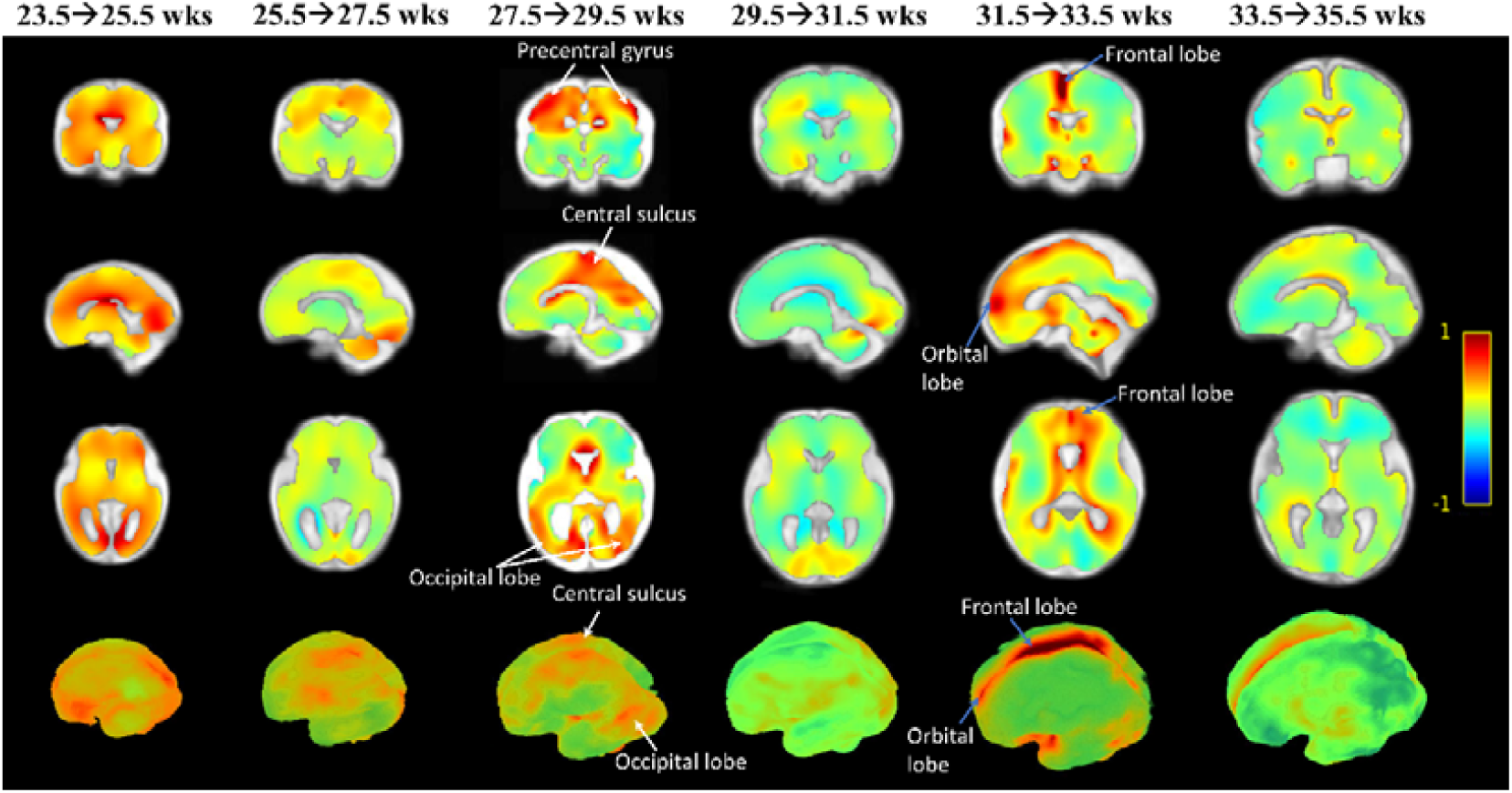
The morphological changes of fetal brains between adjacent gestational stages. The determinant of the log-Jacobian matrix, which represented the amount of morphological change between adjacent fetal brain templates, was rendered in 2D and 3D views. The colors indicate the amount of morphological expansion (red) or retraction (blue, e.g., when cortical folding takes place).

## Discussion

In this work, we proposed a fully automatic segmentation method based on U-net, and together with the SR reconstruction, 3D images of the fetal brains were obtained in an automated manner. Moreover, we generated a 4D spatiotemporal atlas in a Chinese population, based on which, we quantitatively mapped the fetal brain development from 23-36 weeks of gestation. To the best of our knowledge, the existing fetal brain atlases were collected from the Caucasian or mixed populations, which might not be entirely appropriate for analysis of fetal brain development in a different race, as indicated by many studies (Liang et al., 2015; Rao et al., 2017; Tang et al., 2010; T. Zhao et al., 2019). The establishment of a dedicated Chinese fetal brain atlas provided a normative brain template for anatomical reference in clinical diagnosis and quantitative characterization of fetal brain development in related studies.

For quantitative assessment and the volumetric reconstruction of the fetal brain, accurate and automatic fetal brain segmentation is the prerequisite. Extracting the fetal brain from the in utero MRI image is an entirely different task than skull stripping in the adult brain. Due to the complex in utero compositions (amniotic fluid, placental, fetal body), the maternal body tissues surrounding the fetal brain, and the random fetal brain orientation, the fetal brain is often not the gravitational center of the image. Therefore, traditional brain extraction methods that rely on image registration (Taimouri, Gholipour, Velasco-Annis, Estroff, & Warfield, 2015; Tourbier et al., 2017; Wright et al., 2014) mostly fail. Deep-learning methods open a new avenue for this challenging task (Ebner et al., 2019; Khalili et al., 2019; Salehi et al., 2017; L. Zhao et al., 2019). Salehi et al. segmented the fetal brain with two different Auto-Net architectures, including a voxelwise CNN architecture and a fully convolutional network based on the U-net architecture, which achieved Dice scores of 0.9597 and 0.9380, respectively, using a dataset of 75 images (Salehi et al., 2017). Ebner et al. proposed two separate CNNs for localization and segmentation of the fetal brain using 114 scans from normal and spina bifida fetuses for training, and achieved a Dice coefficient around 0.935 (Ebner et al., 2019). Both studies have an optimized CNN structure, but the segmentation accuracies were moderate, possibly due to the limited training data. There were also other studies using deep learning methods for fetal brain tissue segmentation (Khalili et al., 2019; L. Zhao et al., 2019). Here, we proposed a U-net model to segment the fetal brain from 212 scans, which achieved an average Dice of 0.97(±0.01) and robust performance for images in all three orientations across a full gestational age from 23-38 weeks. Considering that the segmentation was performed on routine clinical scans with relatively thick slices and variable image qualities, the segmentation accuracy was comparable or even superior to the existing studies and was sufficient for the subsequent SR reconstruction. Besides, the 2D U-net was computationally efficient, which only took 2-3 seconds to extract one fetal brain. This accurate, robust, and convenient tool is readily translatable to clinical studies.

The generation of fetal brain atlases becomes feasible with the automated fetal brain extraction and SR reconstruction. We took a deformable registration approach to generate a 4D spatiotemporal fetal brain atlas from 23-36 weeks of gestation. A number of population brain atlas generation methods have been proposed (Gousias et al., 2012; Makropoulos et al., 2016; Schuh et al., 2018; Ahmed Serag et al., 2012). Especially, for the generation of the fetal brain atlases, Habas et al. developed age-specific MR templates and tissue probability maps of the fetal brain, based on group-wise registration of manual segmentations and voxel-wise nonlinear modeling (Habas et al., 2010). Gholipour et al. developed an algorithm to construct an unbiased four-dimensional atlas of the developing fetal brain by integrating symmetric diffeomorphic deformable registration in space with kernel regression in age (Gholipour et al., 2017). Here we used iterative SyN registration to ensure gradual convergence of the individual brains to the template. The fact that the iteration converged in 15 iterations indicated the average template became representative of the individual brains at a given GA. This approach has been used in generating neonatal brain atlases (Alexander et al., 2017) and SyN has been demonstrated to be among the best performing deformable registration methods (Klein et al., 2009; Ou, Akbari, Bilello, Da, & Davatzikos, 2014). For a 4D atlas with a temporal dimension, some strategies have been proposed to improve the temporal consistency of atlases between timepoints, such as the adaptive kernel regression method (Ahmed Serag et al., 2012) and the groupwise approach (Schuh et al., 2018). However, since we had a limited number of brains (n=35) and their GA information was limited to integers of weeks, these strategies did not apply to our dataset.

High-quality fetal brain atlases in the Caucasian population have been reported such as (Ahmed Serag et al., 2012; A Serag et al., 2012) and (Gholipour et al., 2014; Gholipour et al., 2017; Khan et al., 2019). Several comparative studies showed considerable anatomical differences between races in children and adults, in terms of the bran size, shape, and topology (Liang et al., 2015; Rao et al., 2017; Tang et al., 2010; T. Zhao et al., 2019). For instance, Zhao et al. showed that in pediatric brains between 6-12 years old, the major anatomical differences between Chinese and Caucasian brain templates were located in the bilateral frontal and parietal areas (T. Zhao et al., 2019). Tang et al. found significant differences in brain shape and size between Chinese and Caucasian young males, such as the brain length, width, height, AC-PC line distance, and their perspective ratios (Tang et al., 2010). It is possible that the developmental differences begin in the fetal period, and therefore, it is essential to build a racial specific fetal brain atlas for related studies. Furthermore, a comparison of the racial specific atlases in the fetal and later stages may help us to understand the genetic versus environmental contributions during brain development.

In addition to visual examination of the fetal brain development from the 4D atlas, we quantitatively characterized the morphological changes between gestational stages based on the deformation maps. For instance, the deformation maps revealed that the primary gyrus, pre- and post-central gyri, and secondary gyrus developed in sequential order. The fast changes in the central sulcus and the precentral/postcentral gyri indicated early development of the preliminary sensory areas, while the subsequent changes in the superior and frontal gyri may relate to the development of higher-order functions. The timeline captured by the deformation maps agreed well with the critical milestones of fetal brain development (Garel et al., 2001; Garel, Chantrel, Elmaleh, Brisse, & Sebag, 2003).

There are several limitations in the current study. First, all of our imaging data were retrospectively collected from pure clinical scans, which typically had low resolution (thick slices), and therefore, the reconstructed 3D images may not match the existing high-resolution Caucasian fetal brain atlas (Gholipour et al., 2017). The low slice resolution limited more detailed analysis, such as GM and WM segmentation. However, they were sufficient for morphological analysis as we have shown, and they were suitable references for analysis of clinical fetal MRI data acquired at similar resolution. Second, we did not perform a quantitative comparison between the Chinese fetal brain atlas with the existing Caucasian atlases in this study due to the unmatched image resolution as explained above. Further work will revisit this scientific question once higher resolution data become available. Moreover, the number of normal developing fetal brain samples was relatively small. Therefore, we were not able to generate a population template for each gestational week but combined data for every two gestational weeks, assuming relatively small anatomical change within the two-week periods. Finer GA intervals should be used when more data become available. Lastly, the U-net method was not directly compared with other neural networks proposed by other groups, as the current segmentation accuracy was sufficient for SR reconstruction.

## Conclusion

In this work, we proposed an automated fetal brain analysis pipeline, including brain extraction, SR reconstruction, atlas generation, and quantification of brain morphological development. The U-net model yielded superior segmentation accuracy compared with the conventional brain extraction method. Using this automated approach, we were able to reconstruct fetal brains across gestation and generate a 4D fetal brain atlas between 23-36 gestational weeks in a Chinese population. The spatiotemporal atlas allowed us to depict normal in utero fetal brain development, which provides normative references for fetal brain examinations in clinical practice.

## Contributions

HL processed MRI data and performed analyses. DW was in charge of the study design and overall progress of the study. HL and DW drafted the manuscript. GY, KL, and YZ contributed to the collection of the MRI data. All authors contributed to the interpretation and reviewing of the manuscript.

## Data and code availability and statement

The code used in this work is available from the authors upon request. The data (anonymized) in this study can be accessed with a data-sharing agreement.

