## Supplementary Figures for "Mapping fetal brain development based on automated segmentation and 4D brain atlasing"

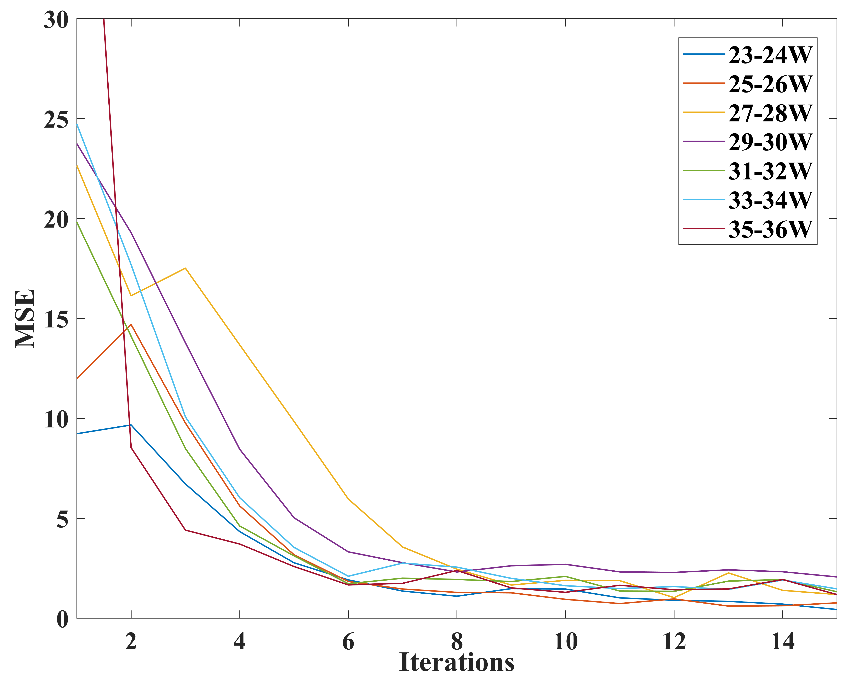

**Figure S1:** Mean square error (MSE) between the averaged brain templates from adjacent iterations during the atlas generation process for different gestational stages.

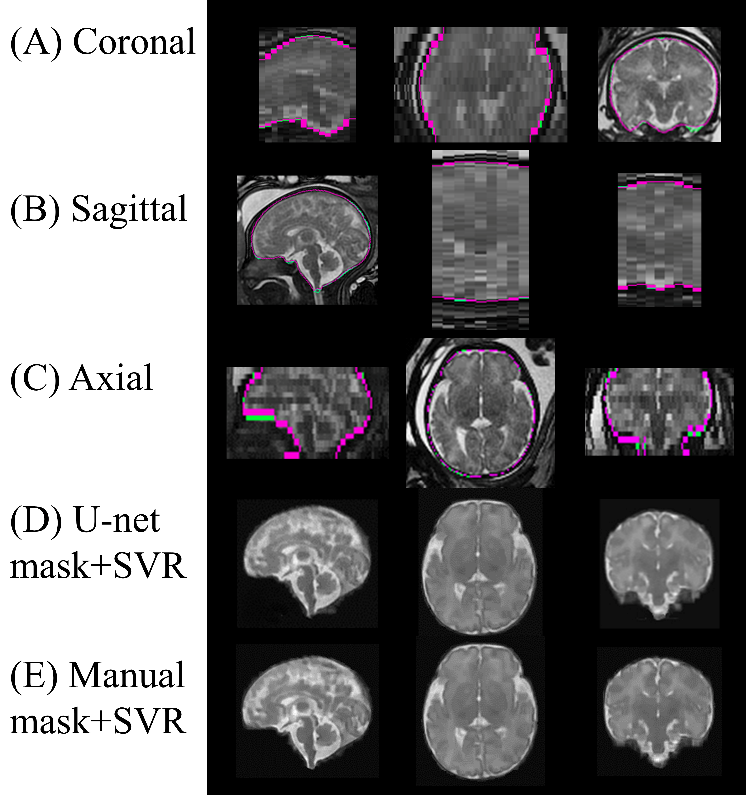

**Figure S2:** An example of the super-resolution reconstruction results based on manually mask and U-net output, in a fetal brain at 34 weeks of gestation. (A-C) The 2D multislice images were shown in the coronal, sagittal, and axial orientations. The contours of the U-net predicted mask are in magenta and the ground truth manual mask are in green. The super-resolution results based on the U-net predicted mask (D) were compared with those using the manually defined brain mask (E).

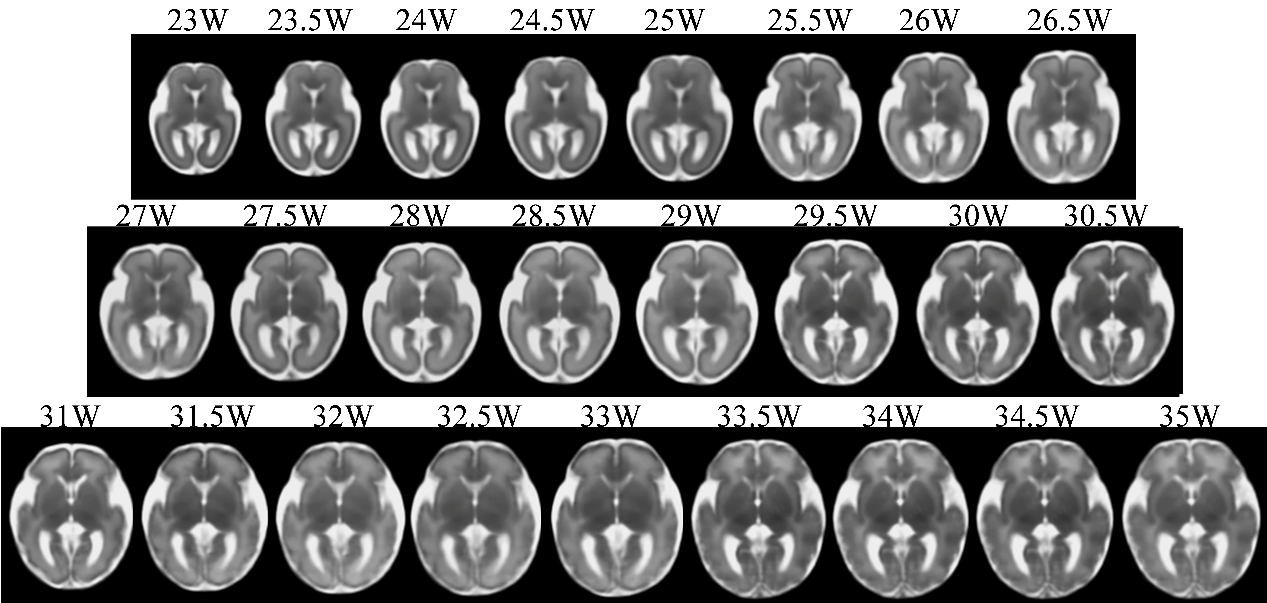

**Figure S3:** Fetal brain templates generated at a 0.5-week interval between 23-35 gestational weeks by interpolating the transformation matrices between the original atlases.

**Supplementary Video**

Dynamic changes of the fetal brain atlas between 23-36 weeks of gestation in a 0.2-week increment.
