## Supplementary figures and images for "Mapping fetal brain development based on automated segmentation and 4D brain atlasing"

### Supplementary Video

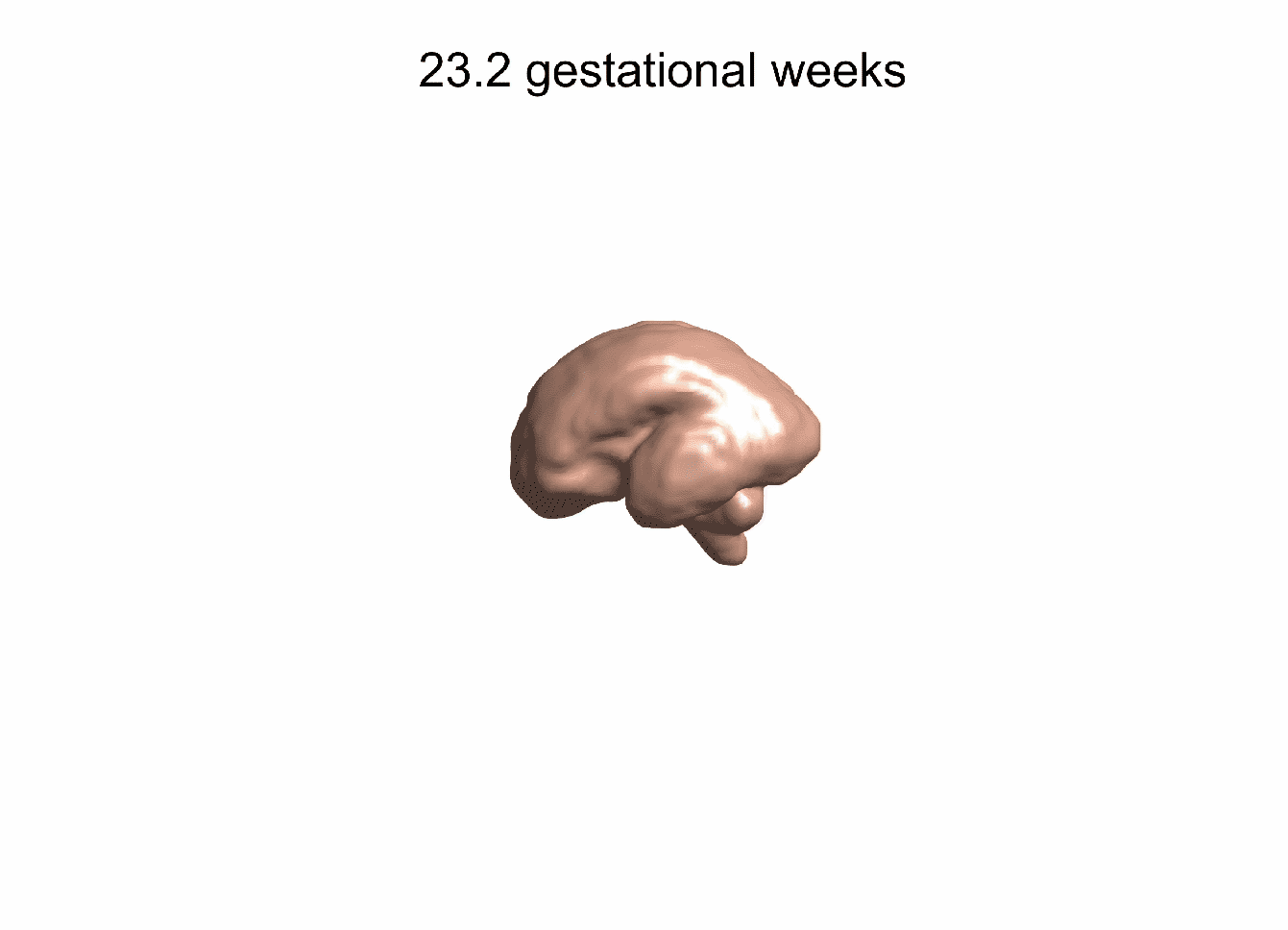
